# Identification and Validation of Small Molecules with Mucin-Selective Regiospecific Binding in the Gastrointestinal Tract

**DOI:** 10.1101/2025.03.31.646052

**Authors:** Deepak A. Subramanian, Ameya R. Kirtane, Georgia N. White, Dylan E. Freitas, Keiko Ishida, Josh Jenkins, Andrew Pettinari, Josh Morimoto, Nina Fitzgerald, Giovanni Traverso

## Abstract

Oral drug delivery is a widely used method of drug administration; however, achieving localized drug release at specific regions of the gastrointestinal (GI) tract is generally accomplished by using broad environmental differences. The GI tract is a complex system with regional differences in composition, such as selective expression of mucin glycoproteins in different organs. Here, we identify small molecule ligands that can selectively bind to the different mucins to localize drug delivery to the small intestine and stomach. We demonstrate up to a 10-fold increase in particle binding to these organs and up to a 4-fold increase in selectivity compared to chitosan. Additionally, we observe up to a 9-fold increase in budesonide concentration in the small intestine and a 25-fold increase in tetracycline concentration in the stomach. These results show that we have developed a versatile platform capable of sequestering a variety of drugs in certain GI tract organs.

## 1. Introduction

Oral drug delivery is commonly used for drug administration because it is less invasive, less painful, and easier to administer when compared to parenteral drug delivery ^[1]^. As such, oral drug delivery may increase treatment efficacy because of greater patient adherence to the drug therapy regimen ^[2]^. However, various barriers of the gastrointestinal (GI) tract inhibit successful drug delivery, such as the hostile luminal environment characterized by the presence of degrading enzymes, bile salts, and food ^[3]^, poor gastrointestinal retention ^[4]^, and poor drug targeting, which can lead to inefficient drug uptake ^[5]^. Developing methods to overcome these barriers is essential to improve drug delivery ^[6]^.

Nanoparticles have been used for oral drug delivery in order to protect the drug cargo from degradation and to control the release profile to optimize drug pharmacokinetics ^[7, 8]^. The materials used for nanoparticle synthesis can be chosen to improve drug delivery by optimizing drug localization and lengthening the drugs’ GI transit time ^[9]^. One method of doing so involves mucoadhesion ^[10]^; in this process, the nanoparticles bind to the mucus layers that cover the epithelial surfaces of the GI tract, allowing for them to remain in one place and release the drug rather than passing through normally ^[11]^. Mucoadhesive approaches have been demonstrated for a variety of different nanoparticles and encapsulated drugs ^[12]^; however, these approaches have generally relied on non-specific binding to mucus utilizing electrostatic, wetting, and mechanical interlocking interactions with the negatively charged mucus ^[11, 13]^. While this does improve residence in the GI tract, it does not allow for targeting due to the ubiquitous presence of mucus across the length of the GI tract. This means that the drug may release in a location that is not optimal for drug absorption. Furthermore, the general mucoadhesive approach is less effective at treating location-specific diseases, such as ulcerative colitis in the small intestine or colon or ulcers in the stomach, because of the relatively poor targeting capability of this approach.

To increase the localization of these therapies to specific parts of the GI tract, we examined the organization of the GI tract and identified regional differences in mucus layer structure. The mucus layers, which serve to cover the epithelial tissue in the GI tract, allow for nutrient diffusion into the epithelium, and prevent the surface from pathogenic attack, are composed of mucin glycoproteins ^[14]^. These glycoproteins are synthesized in goblet cells found throughout the GI epithelium and are secreted to form the mucus layers ^[15]^. The mucin composition of the mucus layers differ in the distinct regions of the GI tract; for example, the mucus layers in the small intestine and colon are composed primarily of the mucin MUC2 ^[16]^, the mucus layers in the stomach are primarily composed of the mucins MUC5AC and MUC6 ^[17]^, and the mucus layers in the salivary glands and esophagus are primarily composed of the mucin MUC5B ^[18]^. While these mucins share many structural similarities (such as D domains, cysteine-rich regions, and “PTS” domains), the chemical structure is distinct for each mucin, presenting unique targeting opportunities. For example, MUC2 is a simpler structure, in which large PTS domains are separated by two cysteine-rich domains; in contrast, MUC5AC is a larger glycoprotein with multiple cysteine-rich regions separating the PTS domains ^[19]^. Aspects such as the greater presence of cysteine-rich regions in MUC5AC compared to MUC2 show a chemical distinction between the two mucins that could potentially be exploited to target one mucin over the other. Developing methods of targeting these mucins would allow for greater localization of drug nanocarriers in addition to the greater residence time conferred through their mucoadhesive properties, allowing for greater control over the region of drug release (**Figure 1**). This would allow for both greater localization of drug absorption in the optimal location for particular drugs, as well as the possibility of treating location-specific diseases by localizing the treatment in the specifically affected region of the GI tract.

**Figure 1.**
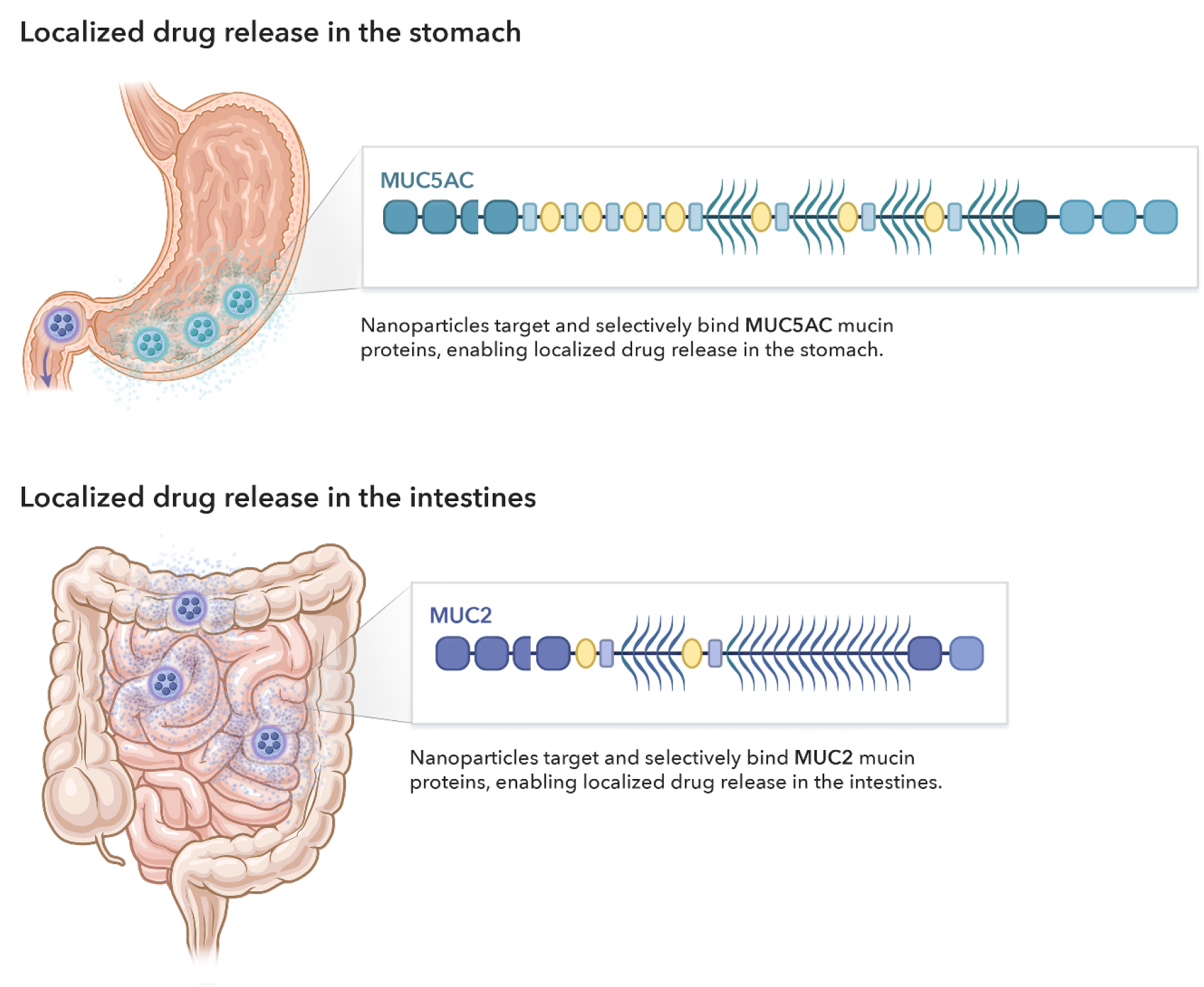
Localized drug delivery approach. Schematic of stomach-specific and intestine-specific regioselective drug delivery approaches based on binding to the mucin glycoproteins found in these environments. Legend on the bottom shows which elements of the structure correspond to different regions of the mucins (D-domain, cysteine-rich region, PTS domain, and cysteine knot). Schematic of mucin structures adapted from Subramanian *et al*. ^[9]^.

While proteins are generally used as ligands to target various regions of the body, their usage in the GI tract can be challenging due to the its harsh environment, which can cause issues such as pH-mediated denaturation and degradation by peptidases and bile salts ^[8]^. While small molecules have been widely used for targeting proteins for therapeutic purposes, they have rarely been used for localization of drug release in the GI tract. One example of their use is the development of folic acid-conjugated nanoparticles for targeting folate receptors in cancer cells ^[20]^. Therefore, small molecules will likely have greater stability in the GI tract than macromolecules. As such, we decided to use small molecules as the ligands of interest in targeting the mucin glycoproteins.

In this work, we identify small molecule ligands that can bind selectively to three different mucins: MUC2, MUC5AC, MUC5B. We first screen a selection of 72 diverse molecules in an *in vitro* screening process to identify those that bind to the mucins more strongly and selectively than chitosan, a known general mucoadhesive. We then validate the performance of these hits in an *ex vivo* setting on freshly dissected porcine tissue to demonstrate their capability to bind to an intact mucus layer. Next, we measure the *in vitro* binding kinetics and thermodynamics of these top hits to confirm the selective nature of binding. Finally, we demonstrate their ability to selectively bind *in vivo* in a rodent model and provide proof of concept for their use in regioselective therapeutic delivery.

## 2. Results

### 2.1 Molecules were selected according to their physicochemical properties to ensure a diverse library

From a selection of 8980 possible molecules, a group of 72 was selected that most clearly represented the diversity of the library; this was based on the physicochemical properties of the molecules in the original set (**Figure 2a-b). Table S1** shows the selection of 72 molecules, along with their index numbers. These numbers are used to represent the molecules in future sections.

**Figure 2.**
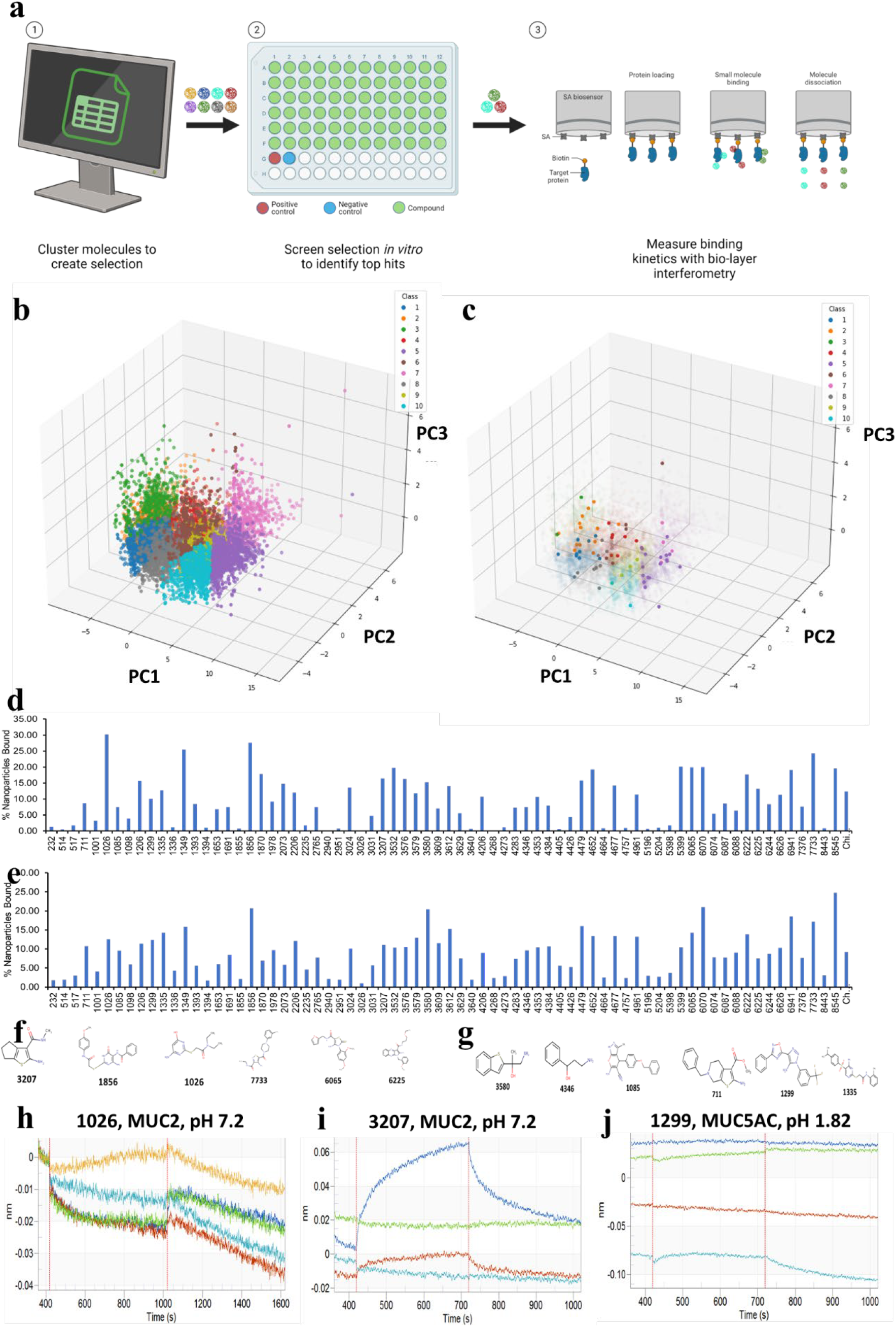
Library selection and *in vitro* binding results. The overall workflow is shown in (a). Principal component analysis and *k*-means clustering was used to cluster the original library of 8980 compounds across three principal components (b). The representative library of 72 compounds was chosen by selecting those nearest to the cluster centroids (c). Results for MUC2 (d) and MUC5AC (e) binding are shown as the median of five replicates. Identified MUC2-selective (f) and MUC5AC-selective (g) hits are shown. Bio-layer interferometry curves for hit 1026 (h), hit 3207 (i), and hit 1299 (j) are shown, with the association (left) and dissociation (right) phases separated by the central red line. The curves shown for hit 1026 and hit 3207 represents binding to MUC2 at pH 7.2, while the curve shown for hit 1299 represents binding to MUC5AC at pH 1.82. Figure 2a created with BioRender.com.

### 2.2 Molecules were successfully conjugated to fluorescent nanoparticles

The 72 molecules were conjugated to fluorescent nanoparticles to create a library of conjugated nanoparticles that could be screened for mucin binding. The effect of conjugation on the nanoparticle properties was characterized by measuring the nanoparticle size and surface zeta potential (using DLS). As compared to unconjugated nanoparticles, which have a nanoparticle size of 100 nm and a zeta potential of -40 mV (due to the negatively charged carboxyl groups), the conjugated nanoparticles showed an increase in nanoparticle size (up to 250 nm) and a zeta potential that was less negative (shown in **Figure S1**).

### 2.3 Molecules were identified that bound selectively to mucins in vitro

The library of conjugated nanoparticles was then incubated with mucin-coated plates. Unbound and weakly bound nanoparticles were washed away, and the remaining strongly bound nanoparticles were dissolved to quantify the amount of fluorescent dye, which could be correlated with the mass of nanoparticles remaining (**Figure 2c**). The percentage of nanoparticles bound (**Figure 2d/e**) and the selectivity for the two mucins (**Figure S2**) were then reported. Binding to MUC5AC was more uniform across most molecules, with some deviations. In contrast, binding to MUC2 varied tremendously; some molecules did not bind at all, while others bound very well. This led to selectivity generally skewing towards MUC5AC, with some molecules binding only to MUC5AC. Because the magnitude and selectivity of binding are both important for any successful mucin-binding molecule, only molecules that showed superior binding and selectivity to MUC2 and MUC5AC compared to the known mucoadhesive, chitosan, were considered as hits. After repetition of the study to confirm the results, six hits (**Figure 2f/g**) were identified for both MUC2 and MUC5AC, which progressed to the *ex vivo* validation stage.

### 2.4 Ex vivo validation identified molecules that could bind selectively to mucin-containing tissues

The six hits were then tested in an *ex vivo* setting (**Figure S3**), where they were allowed to bind to a segment of porcine tissue and washed to remove unbound nanoparticles; fluorescence imaging was used to semi-quantitatively compare nanoparticle binding (**Figure 3a**).

**Figure 3.**
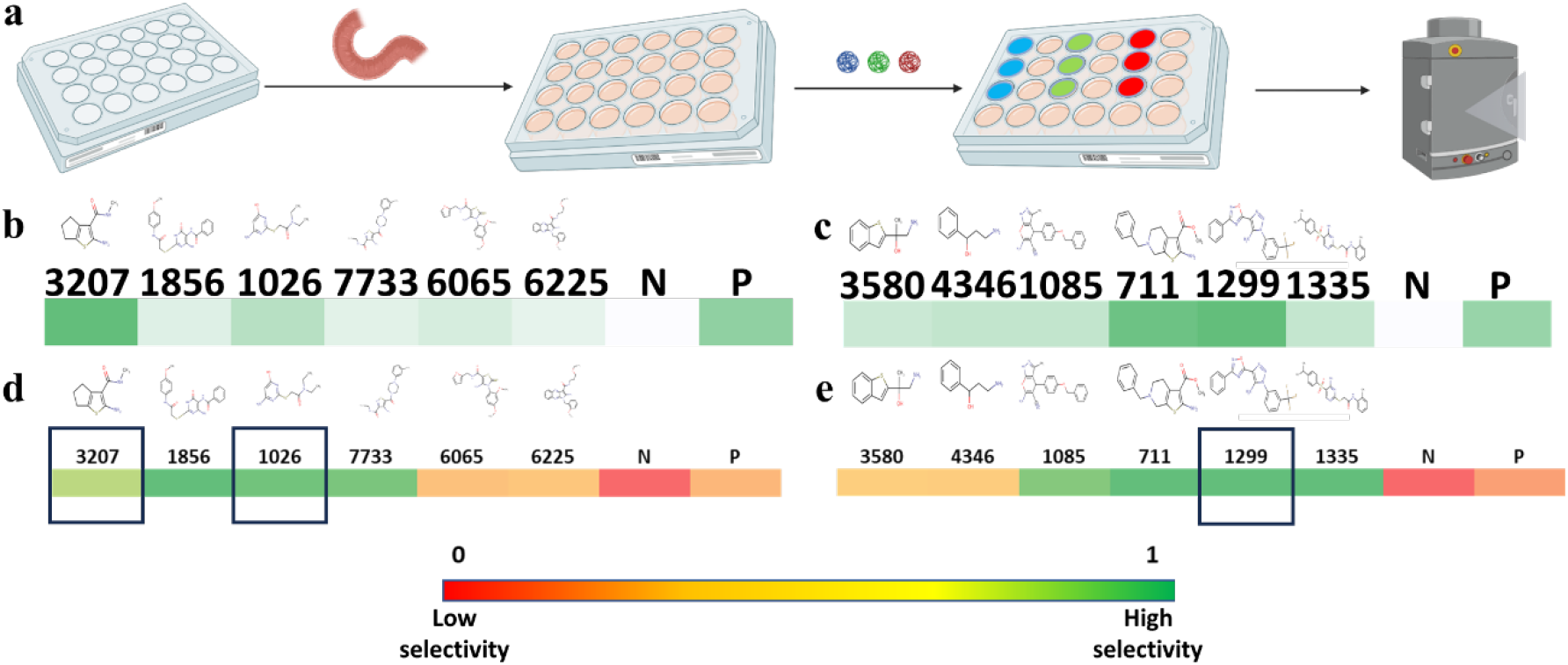
*Ex vivo* screening results. *Ex vivo* screening of the small molecule library and *ex vivo* validation of the identified small molecule “hits” for MUC2 and MUC5AC binding, with a schematic of the screening/validation process (a), the average *ex vitro* binding magnitude for each hit to MUC2 (b) and MUC5AC (c), and the *ex vivo* selectivity results and strongest hits for MUC2 (d) and MUC5AC (e). Darker colors in (b) and (c) represent higher magnitude, while the legend below (d) and (e) shows the relationship between color in the figures and selectivity. Figure 3a created with BioRender.com.

The “score” results for each set of hits across a range of gastrointestinal tissues (small intestine, stomach, and esophagus) are presented (**Figure 3b/c** for mucin of interest, **Figure S4** for other mucins). There was some variability in the results, but the hits generally bound better to the tissues containing their respective mucins. MUC2 hits bound better to the small intestine and MUC5AC hits bound better to the stomach, compared to other gastrointestinal tissues. Based on the “score” results, “selectivity” results were calculated and presented (**Figure 3d/e**). The top hits (3207 and 1026 for MUC2 binding and 1299 for MUC5AC binding) were identified based on both the “score” and “selectivity” results, indicating hits that had both high binding performance and selectivity.

### 2.5 Mucin-binding kinetic affinity of top hits shows strong and selective binding

The mucin-binding kinetic constants and dissociation constant measured from the BLI for the identified hits can be found in **Table S2**. Binding curves for the hits to the mucins of interest at normal pH are shown in **Figure 2h-j**, while the binding curves for off-target mucins and/or on-target mucins at different pH are shown in **Figure S5**. Hit 3207 followed a 2:1 heterogeneous ligand binding model when binding to MUC2 and a 1:1 bimolecular binding model when binding to MUC5AC, while hits 1026 and 1299 followed a 1:1 bimolecular binding model. Hit 3207 showed micromolar binding affinity to MUC2 at neutral pH and nanomolar affinity at pH 3.05, indicating that the molecule bound more tightly to MUC2 at lower pH. When looking at the kinetic rate constants for the forward and reverse reactions at lower pH, we observed faster binding to MUC2, as indicated by the higher rate constant in the forward direction. We also noted a slower desorption from MUC2, as indicated by the lower rate constant in the reverse direction, for hit 3207 when compared to its kinetic behavior at neutral pH. However, when looking at the results for binding to MUC5AC, we saw no improvement in binding affinity, indicating that the 3207 would not be expected to bind selectively to MUC2.

In contrast, we saw that hit 1026 bound more strongly to MUC2 than hit 3207 did at both pH ranges due to its picomolar binding affinity; when looking at the kinetic rate constants, the forward reaction occurred more quickly at acidic pH, indicating faster binding. However, the reverse reaction proceeded more slowly at neutral pH, which indicated slower desorption. Encouragingly, the dissociation constant for the binding of hit 1026 to MUC5AC was 2-3 orders of magnitude larger than its dissociation constant for MUC2, indicating that hit 1026 should be able to bind selectively to MUC2 compared to MUC5AC.

Hit 1299 showed different binding behavior than the other two hits. At neutral pH, it showed relatively weak binding to MUC2. However, at a more acidic pH, the binding affinity became much stronger, similar to that of the 1299-MUC5AC binding interaction at strongly acidic pH. The significant difference was that the picomolar binding affinity of hit 1299 to MUC2 at acidic pH was driven by a very high forward reaction rate constant, indicating that binding occurred relatively quickly. However, desorption also occurred relatively quickly, as evidenced by the relatively high reverse reaction rate constant. This indicates that prolonged retention may be more challenging. In contrast, the picomolar binding affinity of hit 1299 to MUC5AC at stomach pH was driven primarily by a very slow reverse reaction rate constant, indicating that desorption took place much more slowly. This was a promising indicator that hit 1299 could be used for prolonged retention to MUC5AC-containing mucin layers in the stomach, despite its lower mucin-binding selectivity than hit 1026 to MUC2.

### 2.6 Identified hits show localization to mucin-containing tissues in vivo

The performance of the top hits identified from the *in vitro* and *ex vivo* screening (3207 and 1026 for MUC2 small intestinal binding and 1299 for MUC5AC gastric binding) were then validated in a rat model. The first validation step was to ensure that these molecules could bind selectively to the mucus layers of interest, and so we looked at their biodistribution along the GI tract upon oral gavage. We first administered hit-conjugated nanoparticles, along with our controls, to rats via oral gavage. After 8 h, we imaged and analyzed the GI tracts of the rats to identify the location of the nanoparticles. We examined three regions of the GI tract (esophagus, stomach, and intestines) which contained different mucins, to identify both the magnitude and selectivity of binding to the region of interest.

The major results of the small intestinal binding and selectivity for hits 3207 and 1026, the stomach binding and selectivity for hit 1299, and the results for the negative and positive control in each case are shown in **Figure 4a-d**, with more detailed results found in **Figure S6** and **Tables S3-S4**. All results have subtracted the fluorescence from tissue from untreated rats. The hits generally showed a significant increase in total binding throughout the GI tract when compared to the negative control (**Figure 4a-b**). Hit 1026 showed the greatest binding in the small intestine, a 2.5-fold improvement over the positive control, while hit 3207 showed comparable binding magnitude to the positive control. Both hits demonstrated superior binding, an 8-9-fold improvement, compared to the negative control (**Figure 4c, Figure S6d**). While selectivity results are slightly affected by the normal passage of particles through the GI tract, both hits showed high selectivity, with hit 1026 again showing the greatest selectivity (**Figure S6h**). Hit 1299 showed the greatest magnitude of stomach binding when compared to the controls, with a 10-fold improvement compared to the negative control and a 4-fold improvement compared to the positive control (**Figure 4d, Figure S6f**). In addition, hit 1299 showed a 3-fold improvement in stomach-binding selectivity compared to the positive control (**Figure S6i**), demonstrating that hit 1299 can improve localization to the stomach compared to a general mucoadhesive.

**Figure 4.**
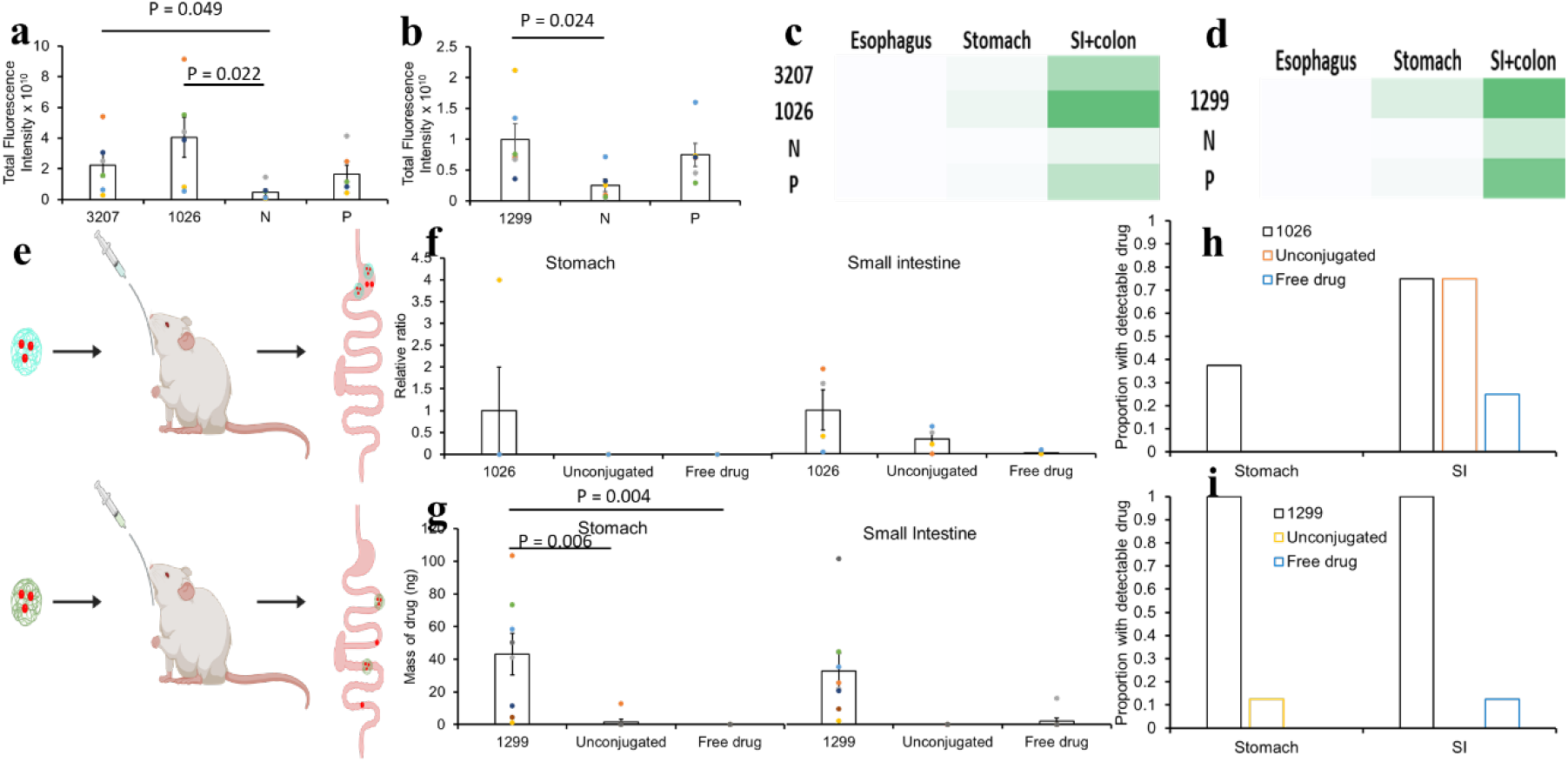
*In vivo* validation results for MUC2-selective and MUC5AC-selective hits. First, the biodistribution was observed. The total binding fluorescence for MUC2-selective (a) and MUC5AC-selective (b) hits was measured across the entire GI tract, and the results for each organ are shown as heat maps for the MUC2-selective (c) and MUC5AC-selective (d) hits, where darker colors represent greater binding. The schematic for the proof-of-concept for localized drug delivery is shown in (e). The budesonide concentration results for the different MUC2-binding formulations are shown by comparing the relative ratio of drug (compared to the MUC2-selective hit) for each formulation at all locations (f). Tetracycline results are shown by comparing the amount of drug present at all locations for the hit and controls (g). Error bars represent standard error of measurement (n=6 for 4a-d, n=4 for 4f, n=8 for 4g).

### 2.7 Top hits can be used for tissue-localized drug delivery

The top performing hits (hit 1026 for MUC2-selective binding and hit 1299 for MUC5AC-selective binding) were then conjugated to drug-loaded PLGA-*b*-PEG nanoparticles and used to demonstrate their capability for localized drug delivery. **Figure 4e** shows the overall schematic for this process. The major results are shown in **Figure 4f-g** with all data shown in **Tables S5-S6**. When comparing the relative amount of budesonide in the small intestine across formulations, the hit-conjugated nanoparticles showed a 3-fold increase in budesonide compared to unconjugated nanoparticles and a 50-fold increase in budesonide compared to a suspension of free drug (**Figure 4f**), indicating that the residence of budesonide was prolonged due to enhanced binding of the formulations to MUC2 in the small intestine. For stomach targeting, we saw a significantly higher concentration of tetracycline in the stomach for the hit-conjugated nanoparticles compared to the other two formulations, with a 25-fold increase in tetracycline concentration compared to the unconjugated nanoparticles (**Figure 4g**). This validates the performance of the hits in localizing drug to the stomach and increasing drug residence.

## 3. Discussion

Oral drug delivery, while the most widely used method of drug administration, has many challenges that affect the efficacy of drug uptake. It is particularly critical to ensure that the drug is released at the site of absorption long enough for a clinically relevant concentration to be achieved within the therapeutic window ^[21]^. To address this, we identified and validated the performance of small molecules that can bind strongly and selectively to secreted gastrointestinal mucin glycoproteins (MUC2, MUC5AC), which compose the mucus barriers within the GI tract. Because these mucins are only expressed in certain regions of the GI tract (MUC2 in the small intestine/colon and MUC5AC in the stomach), we hypothesized that a regioselective mucoadhesion approach could improve both the retention time and localization for oral drug delivery systems.

We first screened an initial selection of molecules to identify those that bound strongly and selectively to the mucins *in vitro* and *ex vivo*. We then investigated the strength of binding of the successful candidates by measuring the binding kinetics and calculating the dissociation constant, demonstrating that the most promising molecules bound both strongly (nanomolar affinity) and selectively (reduction in dissociation constant by at least one order of magnitude) to the mucin of interest. Finally, we showed that the molecules could aid localization and retention of various payloads (fluorescent dye, budesonide, and tetracycline) in a rat model. The results shown here indicate that conjugating these small molecules to the surface of a payload (such as a nanoparticle or drug) can improve their localization and retention in a particular region of the GI tract when delivered orally. This is an attractive option for a couple of different reasons. First, this targeting can provide more effective treatment of local GI diseases by ensuring that the drug is mostly released at the site of the disease. For example, this could aid treatment of stomach ailments caused by *H. pylori* infection, as most of the antibiotic could be delivered to the stomach, rather than losing some of the drug to release and absorption in the small intestine. This could lead to the development of shorter-term drug regimens while maintaining the efficacy of treatment, which could reduce the chance of antibiotic resistance ^[22]^. In addition, providing strong mucoadhesion would result in increased residence within the GI tract, which could improve the capability of large-dose drug delivery. Finally, it can allow for selective drug release at specific sites of the GI tract, which may aid in the absorption of select drugs. For example, the drug furesomide has been shown to be preferentially absorbed in the stomach, with Klausner *et al*. (2003) indicating that developing a gastroretentive formulation could increase the length of the absorption phase of furesomide in the stomach 3-fold ^[23]^. Other drugs are poorly absorbed in the stomach and show higher bioavailability when absorbed in the small intestine or colon; for example, Sampathi *et al*. (2023) demonstrated that preparing curcumin nanosuspensions that allow for colonic delivery can improve the bioavailability 4-fold ^[24]^.

Two major limitations of the work that can be expanded upon are the limited diversity of screened molecules and the relatively wide range of target. First, increasing screening throughput could allow for testing of more diverse molecules. Due to the employed method of screening, we tested 72 molecules due to the need to individually conjugate these molecules to the surface of fluorescent nanoparticles. We also limited our search to primary amines due to the ease of conjugation to carboxyl-functionalized nanoparticles. However, improvements in high-throughput screening technology such as DNA-encoded libraries ^[25]^ and small molecule microarrays ^[26]^ can be used to screen thousands of molecules simultaneously against targets of interest such as the mucin glycoproteins, which can greatly increase the diversity of the chemical space that could be screened. Furthermore, increasing the size of the generated dataset can allow for the use of more complex AI models such as graph neural networks to identify structural elements of the molecules that are correlated with stronger and more regioselective binding ^[27]^.

Additionally, gastrointestinal targets of increasing specificity can be screened to further refine the regioselectivity of the approach. Our approach involved using differences in mucin composition to promote regioselectivity because this was a relatively straightforward way to localize drug delivery to the different organs, such as the stomach and small intestine, within the GI tract. However, targeting specificity can be further improved by identifying targets within these regions that are expressed differently. For example, the intestinal epithelium contains a variety of different cell types (such as enterocytes, goblet cells, Paneth cells, Peyer’s patches, dendritic cells, proliferating stem cells, neuroendocrine cells) with their own unique functions ^[28]^, and developing methods of targeting these cell types can lead to a greater ability to control their specific functions without affecting other nearby cell types ^[29]^. In addition, the glycosylation of the mucins themselves can be altered in various disease states ^[30]^. Identifying molecules that can target these changes in glycosylation can be used to further improve targeting towards diseased mucus. This can result in increased drug release at the site of the disease, thereby increasing efficacy and reducing side effects due to off-target release.

In summary, we have identified and validated molecules that strongly and selectively bind to the MUC2 and MUC5AC glycoproteins, which are selectively expressed in the intestines and colon, respectively, in the human GI tract. We have additionally demonstrated that these molecules can aid in localizing and retaining nanocarriers containing antibiotic and anti-inflammatory drugs within these regions to improve oral drug delivery. Since these molecules are conjugated to the surface, they can be used to develop a variety of localized therapies that can be employed for various gastrointestinal diseases.

## 4. Materials and Methods

### 4.1 Materials

Small molecules were purchased from Life Chemicals, Inc. (Niagara-on-the-Lake, Ontario, Canada). 1-ethyl-3-(3-dimethylaminopropyl)-carbodiimide (EDC), N-hydroxysuccinimide (NHS), carboxyl-terminated poly(lactic acid-*co*-glycolic acid)-*block*-poly(ethylene glycol) (PLGA-*b*-PEG-COOH), and tetracycline were purchased from Sigma-Aldrich (St. Louis, MO, USA). Carboxyl-functionalized red fluorescent nanoparticles (0.1 µm) and demeclocycline were purchased from Invitrogen (Carlsbad, CA, USA). MUC2 and MUC5AC purified proteins were purchased from MyBioSource (San Diego, CA, USA) and MilliporeSigma (Burlington, MA, USA). Porcine tissue was procured from the Blood Farm Slaughterhouse (West Groton, MA, USA). Budesonide was purchased from Selleck Chemicals (Houston, TX, USA).

### 4.2 Selection of small molecules for screening

Molecules were selected from an original set of 8980 primary amines (Life Chemicals, Inc.), chosen because amines have a wide variety of conjugation chemistries that can be used to couple them to the surface of nanoparticles. To arrive at an initial selection of 72 molecules, JChem was used to compute a selection of 20 physicochemical properties for each of the molecules (JChem version 21.13.0, ChemAxon). Afterwards, principal component analysis (PCA) was used to reduce the dimensionality to a set of 3 principal components. These principal components were then fed into a k-means clustering algorithm to categorize the molecules into 10 clusters. Finally, the clusters were plotted, and a subset of molecules closest to the centroid of each cluster were selected to best represent the cluster. These molecules were purchased and used in subsequent screening.

### 4.3 Small molecule conjugation to fluorescent nanoparticles

The binding of the selected molecules to mucins *in vitro* was measured using a microtiter plate fluorescence-based assay. The candidate molecules (as well as the positive control, chitosan) were first conjugated to the surface of carboxyl-functionalized red fluorescent nanoparticles. Briefly, nanoparticles (10 mg, 500 µL, 20 mg/mL) were filtered with a centrifugal filter (10 min, 4000 rpm) to remove their original solvent, and were resuspended in 1 mL of methanol; probe sonication was used (45% output, 3 cycles of 30 s on and 5 s off) was used to disrupt any particle aggregation. EDC (0.44 mg, dissolved in methanol at 10 mg/ml) and NHS (0.55 mg, dissolved in methanol at 10 mg/ml) were added to this suspension, and allowed to react for 30 minutes at room temperature, with gentle agitation on an orbital shaker. Residual EDC and NHS were removed through another round of centrifugal filtration, and the activated carboxyl nanoparticles were resuspended in methanol (500 µL in a 1.5 mL microcentrifuge tube) and sonicated similarly. To this, a solution of the small molecule (0.398 µmol) dissolved in dimethyl sulfoxide (DMSO, 26.2 µL), was added to the suspension, and the conjugation was allowed to take place for 24 h at room temperature with gentle rotation of the tube. Afterwards, the solution was filtered with a centrifugal filter as before, and the supernatant was retained for analysis. The remaining conjugated nanoparticles were suspended in water, sonicated to disrupt aggregate formation, and filtered again; the wash was retained as well. The remaining nanoparticles were finally resuspended in water, sonicated, and stored at 4°C.

Small molecule-conjugated nanoparticles were characterized for their size and zeta potential using dynamic light scattering (DLS). Size and zeta potential measurements were made using a Malvern Zetasizer. In each case, the mean value was recorded as the average of three measurements. The concentration of the conjugated nanoparticles was calculated through fluorescence measurements and comparison to a standard curve.

### 4.4 In vitro mucin-binding screen

Each screen (n=5) was performed in 96-well U-bottom polypropylene plates (Grenier), in which 74 wells of each plate were used (one for each of the 72 molecules in the screen, plus one well each for non-conjugated fluorescent nanoparticles as a negative control and chitosan-conjugated fluorescent nanoparticles as a positive control). The plates were first incubated with mucin (10 µg per well) dissolved in phosphate-buffered saline (PBS, 150 µL per well). The plates were left overnight in a humidified, sealed container at 4°C with agitation at 50 rpm, allowing the mucin to coat the plastic via nonspecific hydrophobic interactions. The next day, the excess coating solution was removed by slapping the plate dry on a clean paper towel. The plates were then blocked by adding a 1x PBS + 5 mg/mL bovine serum albumin solution (200 µL) to each well and incubating for 1 h at 4°C. After blocking, the plates were washed 3x with 1x tris-buffered saline (TBS) + 0.5% Tween-20 (200 µL). Following this, the plates were used for the binding assay.

The conjugated nanoparticles were sonicated as before (45% output, 3 cycles of 30 s on and 5 s off) to disrupt aggregation. Eighty micrograms of each set of nanoparticles were added to each well, along with the negative and positive controls; they were allowed to incubate with the coated and blocked mucin for 1 h at room temperature under gentle agitation (50 rpm). Before incubation began, the fluorescence of each well was measured (excitation/emission wavelengths were 580 nm/605 nm, respectively), and the absolute mass of nanoparticles in each well was determined via comparison with a standard curve. After the incubation period, the non-binding nanoparticles were removed by slapping the plate on a clean paper towel, and the plates were washed 3x with 1x TBS + 0.5% Tween-20 (200 µL) to remove weakly binding nanoparticles. Finally, the remaining bound nanoparticles were dissolved by adding of tetrahydrofuran (200 µL), and the encapsulated fluorescent dye in each well was measured using a similar fluorescence measurement; the mass of nanoparticles was then calculated with a similar standard curve. The mass of bound nanoparticles was compared to the starting mass of nanoparticles in each well, and the percentage of nanoparticles bound was calculated and reported. In addition, the selectivity of each molecule was calculated by taking the proportion of the total percentage bound values for each mucin. The results reported represent the median of the five values.

### 4.5 Ex vivo validation of successful hits

The top performing hits from the *in vitro* screen (those that showed a higher percentage of particles bound and better selectivity to the mucin than chitosan, a known general mucoadhesive) were tested on *ex vivo* segments of porcine mucin-containing tissues (jejunal, stomach, and esophageal tissue) obtained from local abattoirs or from laboratory-maintained pigs. Tissue was used either immediately after acquisition or after storage at -80°C; tissue was used at most 1 week after acquisition and storage at -80°C.

After acquisition and equilibration to room temperature, the tissue was washed twice with cold 1x PBS (25 mL each) to remove undigested food and other debris. After washing and cleaning, the tissue was mounted between two magnetic plates (with holes corresponding to a 48-well plate) as previously described ^[31]^. Briefly, the tissue was first dissected to fit the dimensions of the magnetic plate setup, to minimize the tissue impedance of the magnetic sealing system for the plate setup. For intestinal tissue, the tissue segments were dissected longitudinally before cutting, while processing the stomach tissue involved cutting a properly sized tissue segment and removing the bottom layers of the tissue (the lamina propria and the muscularis mucosae). After this, the remaining tissue segment was clamped between the two magnetic plates, in such a way that the luminal side of the tissue was facing upwards (as depicted in **Figure S3**). This created a series of “wells” in which the *ex vivo* testing could take place.

After the tissue was mounted, conjugated nanoparticles (40 µg), along with the negative and positive controls, were suspended in 1x PBS (200 µL) and added to the wells created by the magnetic plates; this was done 3x for each formulation to perform the experiment in triplicate. Additionally, plain 1x PBS was added to one well to serve as the background measurement. After the nanoparticle dispersions were added, the plate setup was covered with aluminum foil and allowed to incubate for 1 h at room temperature with gentle agitation (50 rpm). After this point, the nonbinding nanoparticles were removed by slapping the plate setup dry on a clean paper towel, and the setup was washed 3x by adding 1x PBS (200 µL) to each well and gently agitating the setup for 30 s before removing the washing solution. After washing was complete, fluorescence imaging was performed using an IVIS Spectrum fluorescent imaging system (Perkin Elmer), with an automatically calculated exposure time (usually ∼1 s), a 20 cm x 20 cm field of view, and an excitation/emission of 570 nm/620 nm, respectively. The imaging results were analyzed using the Living Image software to calculate the total fluorescence intensity for each well, and the results for each hit or control were averaged across different wells.

To better account for tissue-to-tissue variability, the results for each trial were reported as scores rather than using the raw fluorescence intensity. For each trial, the calculated fluorescence intensities were scaled between the negative and positive controls as follows:

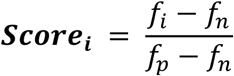

**Equation 1**. Scoring equation for *ex vivo* tissue binding experiments.

Here, *score*_*i*_ represents the hit score for hit *i* (that is reported), *f*_*i*_ represents the mean fluorescence intensity for the hit *i*, and *f*_*n*_ and *f*_*p*_ represent the mean fluorescence intensities for the negative and positive controls, respectively. As such, the score for the negative control will always be 0 and the score for the positive control will always be 1, while the score for the “hits” will be closer to (or greater than) 1 for successful hits and closer to 0 for unsuccessful hits.

The selectivity of a hit *i* to a particular mucin *m* is calculated as follows:

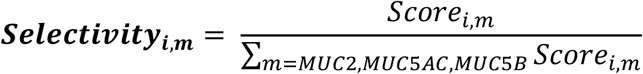

**Equation 2**. Calculation of mucin-binding selectivity for *ex vivo* tissue binding experiments.

Here, the scores shown represent the mean hit score across multiple trials (n=3) for a particular mucin-containing tissue.

### 4.6 Measurement of in vitro binding kinetics

Binding kinetics for the successful mucin-binding hits were measured using bio-layer interferometry (BLI) ^[32]^. In BLI, a protein target is immobilized on the surface of a biosensor, creating a thin optical layer. These coated sensors are then exposed to a solution of the ligand (in this case, the small molecule hits), allowing the ligand to associate and dissociate with the protein target. This association/dissociation behavior changes the thickness of the optical layer, and the instrument correlates this change in thickness with binding and desorption. By testing different concentrations of ligand, the kinetic rate constants and dissociation constant (k_d_) can be measured by fitting the resulting binding curves according to a binding model ^[32]^.

For analysis of mucin-binding kinetics, mucins MUC2 and MUC5AC were used as the protein targets and the “hits” were used as the ligands of interest. Mucins were biotinylated and suspended in buffers with different pH (MUC2 in PBS with pH 7.2 and PBS-HCl with pH 3.35, MUC5AC in PBS-HCl with pH 1.82) that mimic the pH of the normal small intestine ^[33]^, small intestine during inflammatory bowel disease ^[34]^, and the normal stomach ^[35]^, respectively. Hits were dissolved in the same buffer (at concentrations of 100 nM, 200 nM, 400 nM, and 800 nM) to minimize the optical difference between buffer, mucin solution, and analyte solution. BLI was carried out in an Octet RED96 platform with five steps. First, a baseline step of 60 s was carried out. Then, biotinylated mucins were immobilized onto the surface of a streptavidin sensor in a loading step of 300 s. Next, the loaded tips were allowed to re-equilibrate in another baseline step for 60 s. Following that, the loaded tips were allowed to incubate in a solution of analyte (an association step) for 300 s; this allowed the hits to interact with the mucin, changing the thickness of the optical layer and generating a response from the optical measurement. Finally, the tips were moved to buffer wells, and the hits were allowed to dissociate from the mucin for 300 s. Data was analyzed, and ligand-only and hit-only controls were used as references to properly characterize the binding performance of the hits.

Binding of the hits to the mucins generally follows one of two different mechanisms. The first is a 1:1 bimolecular model, in which the molecule binds to a specific spot on the mucin. The reaction can therefore be described using the following model ^[36]^:

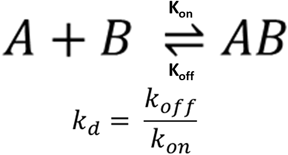

**Equation 3**. Rate law for 1:1 bimolecular binding model.

In this model, k_on_ represents the rate of association of the hit to the mucin, while k_off_ represents the rate of dissociation of the molecule from the molecule-mucin complex. The dissociation constant is calculated as the ratio of these two rate constants.

The second model is a 2:1 heterogeneous model, in which the molecule binds at two locations on the mucin (each with its own kinetic rate constants and dissociation constant) ^[37]^. These reactions are described using the following model:

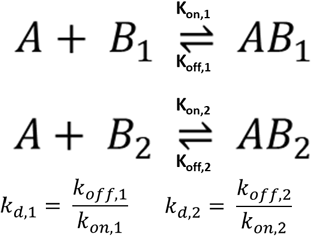

**Equation 4**. Rate law for 2:1 heterogeneous ligand binding model.

The rate constants for each interaction are equivalent to those for the single interaction present in the 1:1 model, while each interaction has its own dissociation constant. This was calculated similarly to that of the 1:1 model.

### 4.7 Gastrointestinal localization of hits in rats

All animal studies were carried out under the supervision of MIT’s Division of Comparative Medicine and in compliance with NIH’s *Guide for the Care and Use of Laboratory Animals*, following the guidelines set forth by MIT’s Committee on Animal Care. Sprague-Dawley rats (200-300 g, mix of male and female) were obtained from Charles River Laboratories and allowed to acclimatize to their new conditions for a week, following which their diet was adjusted to an alfalfa-free diet (AIN-93M, TestDiet) for at least 7 days prior to the study in order to reduce background fluorescence. On the day of the study, rats (n=6) were randomly allocated to the different groups and administered formulations (hits, negative control, positive control, blank) via oral gavage. For rats dosed with the hits and controls, nanoparticles (250 µg) were dosed in 1x PBS (400 µL, 1.25 mg/kg), while the blank was administered PBS only (400 µL). After 8 h, the rats were euthanized through CO_2_ exposure and asphyxiation. The GI tract was extracted (intact) and placed in 1x PBS (50 mL) to remove excess blood from the outside of the GI tract. Fluorescence imaging was performed on the extracted GI tracts using the same IVIS system as before, with an automatically calculated exposure time (ranging between 1-15 s depending on fluorescent intensity), a 23 cm x 23 cm field of view, and an excitation/emission of 570 nm/620 nm, respectively. Imaging results were obtained for the esophagus, stomach, small intestine, and colon, and these were analyzed using the Living Image software. The fluorescence values for the hits and controls were defined as the total fluorescence intensity for the segment of tissue, with the average fluorescence value of the blank for that segment subtracted (to reduce the effect of background autofluorescence), while the selectivity for that segment was calculated as the fluorescence value divided by the sum of the fluorescence values across all three segments of tissue (esophagus, stomach, and intestine).

### 4.8 Demonstration of targeted drug delivery in vivo

PLGA-*b*-PEG-COOH nanoparticles containing encapsulated drugs of interest (budesonide and tetracycline) were prepared via nanoprecipitation in DMF and water as previously described ^[38]^. Budesonide was chosen as a model drug for small intestinal targeting due to its usage as an anti-inflammatory treatment ^[39]^, while tetracycline was chosen as a model drug for stomach targeting due to its usage in the treatment of *H. pylori* infections ^[40]^. Briefly, PLGA-*b*-PEG-COOH and drug were dissolved in DMF (10 mg/mL and 0.1 mg/mL, respectively, in 20 mL DMF), and the resulting solution was added dropwise to a 2x volume of water (stirred continuously at 300 rpm). The resulting dispersion was stirred for 6 h at room temperature. Afterwards, nanoparticles were purified by microcentrifugation (10,000 x *g*, 10 min), washed with water, and centrifuged again. Successful nanoparticle formation was confirmed with DLS particle size measurement and zeta potential measurement, as described earlier; drug loading was measured by dissolving a known volume of nanoparticles in acetonitrile and measuring the drug concentration of the resulting solution using HPLC. Afterwards, the hits were conjugated to the surface of the PLGA-*b*-PEG-COOH nanoparticles using EDC/NHS chemistry (as described earlier). No drug release was noted upon conjugation when performing the conjugation with nanoparticles containing a model dye (Alexa Fluor 647).

Rats were divided into three groups (n=4/group for budesonide and n=8/group for tetracycline): conjugated nanoparticles (C), unconjugated nanoparticles (U), and free drug solution (D). Each rat was dosed drug (20 µg, either encapsulated in the nanoparticles or in suspension) in 1x PBS (500 µL, 0.1 mg/kg) via oral gavage. At a given time point (6 h for tetracycline delivery to the stomach and 12 h for budesonide delivery to the small intestine), the rats were euthanized and the GI tract was harvested. Segments of the stomach and small intestine were selected for homogenization; the stomach was sectioned into three parts, while segments of the duodenum, jejunum, and ileum were selected (totaling 20% of the total mass of the small intestine) due to the size of the rat small intestine and processing limitations. Each tissue segment was homogenized in 500 µL deionized water using a Precellys 24 tissue homogenizer (3 cycles at 4000 rpm), and the resulting homogenate was lyophilized for 72 h. Acetonitrile (1 mL) was then added to the lyophilized powder and mixed overnight at 4°C with gentle agitation (80 rpm) to extract and dissolve any drug and nanoparticles present. The acetonitrile solution containing extracted drug was purified by centrifugation (18,000 x *g*, 10 min) to remove any tissue artifacts.

Tissue samples were analyzed for budesonide using an Agilent 1260 Infinity HPLC coupled to a 6545B quadrupole-time-of-flight (Q-TOF) mass spectrometer (Agilent Technologies, Santa Clara, CA). Samples were injected at a volume of 5 µL onto a Poroshell 120 EC-C18 column (3.0 × 50 mm, 2.7 µm d_p_) held at 40 °C. The mobile phase was pumped at 0.750 mL/min using an isocratic composition of 30% water with 0.1% formic acid (v/v) and 70% acetonitrile over 6 minutes. Agilent JetStream (AJS) electrospray ionization source (ESI) parameters were as follows: gas temperature, 275 °C; gas flow, 10 L/min; nebulizer pressure, 35 psi; sheath gas temperature, 350 °C; sheath gas flow, 12 L/min; capillary voltage, 3500 V; nozzle voltage, 100 V; fragmentor voltage, 110 V. A reference mass mixture was also used to automatically calibrate the mass axis from 121.0509 to 992.0098 *m/z*.

Tissue samples were analyzed for tetracycline using an Agilent 1260 Infinity HPLC coupled to a 6495A triple quadrupole mass spectrometer (Agilent Technologies, Santa Clara, CA). Samples were injected at a volume of 1 µL onto an SB-C18 column (2.1 × 50 mm, 1.8 µm d_p_) held at 24 °C. The mobile phase consisted of 0.1% formic acid in water (v/v, A) and acetonitrile (B), pumped at 400 µL/min with a gradient program of: 0 min, 5% B; 4.5 min, 32% B; 4.6 min, 95% B, with a run time of 6 min and an equilibration time of 1 min. Agilent JetStream (AJS) electrospray ionization source parameters were as follows: gas temperature, 200 °C; gas flow, 20 L/min; nebulizer pressure, 40 psi; sheath gas temperature, 400 °C; sheath gas flow, 12 L/min; capillary voltage, 4500 V; nozzle voltage, 2000 V; high pressure radiofrequency (RF), 110V; low pressure RF, 60V. The collision-induced transition from the [M+H]^+^ ion at 445 *m/z* to 410 *m/z* at 20 V collision energy was used to quantify tetracycline, with the 445 to 427 *m/z* transition at 13 V for signal qualification. Demeclocycline internal standard was quantified with the 465 to 448 *m/z* transition at 19 V and qualified with the 467 to 450 *m/z* transition the same collision energy. The lower limit of quantitation was 1 ng/mL tetracycline in tissue extract.

### 4.9 Statistical Analysis

To determine whether differences between two groups were statistically significant, a two-tailed Student’s *t* test was used. One-way analysis of variance (ANOVA) and post hoc Bonferroni correction were used for multiple comparisons. A *p*-value of less than 0.05 was considered statistically significant.

## Supplemental Information

**Figure S1.** Formulation size (a) and zeta potential (b) for the small molecule-conjugated fluorescent nanoparticles. Baseline values for the nanoparticle size and zeta potential for unconjugated nanoparticles are shown with the red lines.

**Figure S2.** Detailed results from the *in vitro* screening of the 72 hits, detailing selectivity to MUC2 (a) and selectivity to MUC5AC (b). Selectivity is calculated by taking the relative contribution of percentage of nanoparticles bound (seen in **Figure 2**) across all mucins. Results represent the median of five replicates.

**Figure S3.** Mounted setup for *ex vivo* tissue binding. The setup consists of four parts. The bottom plate is a black polystyrene plate (20 cm x 20 cm), which serves to minimize background fluorescence during imaging studies. The next two plates are magnetic plates with 48 wells cut into the plates. These can suspend a piece of tissue through the magnetic force that holds the plates together, with the luminal side of the tissue facing upwards, allowing for incubation of mucin-binding formulations. The final plate is a standard 48-well plate cover, which is used to enclose the tissue setup.

**Figure S4.** Detailed *ex vivo* binding results for MUC2-selective hits to the small intestine (a), stomach (b), and esophagus (c) and MUC5AC-selective hits to the small intestine (d), stomach (e), and esophagus (f). Colored dots represent individual data points. Values on the y-axis correspond to the nanoparticle score, which is defined as 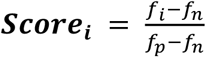. Error bars represent standard deviation. *** - p < 0.005

**Figure S5.** Association and dissociation curves measured by bio-layer interferometry. Results shown are for hit 1026 to MUC2 at pH 7.2 (a), MUC2 at pH 3.35 (b), and MUC5AC at pH 1.82 (c), hit 3207 to MUC2 at pH 7.2 (d), MUC2 at pH 3.35 (e), and MUC5AC at pH 1.82 (f), and hit 1299 to MUC2 at pH 7.2 (g), MUC2 at pH 3.35 (h), and MUC5AC at pH 1.82 (i).

**Figure S6.** Detailed *in vivo* biodistribution results, showing the MUC2-binding hit fluorescence in the esophagus (a), MUC2-binding hit fluorescence in the stomach (b), MUC2-binding hit fluorescence in the small intestine/colon (c), MUC5AC-binding hit fluorescence in the esophagus (d), MUC5AC-binding hit fluorescence in the stomach (e), and MUC5AC-binding hit fluorescence in the small intestine/colon (f). Detailed selectivity results are shown for MUC2-selective (g) and MUC5AC-selective (h) hits. Error bars represent standard error of measurement. **** - *p* < 0.0001.

## Acknowledgments

We are grateful to the Langer and Traverso laboratories for their helpful feedback on the experiments and suggestions for advancing the project. We are grateful to David Cawston for his support in training and assisting researchers in rodent handling and oral gavage techniques. We thank the Koch Institute’s Robert A. Swanson (1969) Biotechnology Center for technical support, specifically the Preclinical Imaging and Testing core facility. V. E. Fulford helped sketch Figure 1.

D.A.S. was sponsored by the NSF Graduate Research Fellowship. G.T. was sponsored in part by the Department of Mechanical Engineering, Massachusetts Institute of Technology and the Division of Gastroenterology, Brigham and Women’s Hospital, Harvard Medical School. This work was sponsored in part by the Koch Institute Support (core) Grant P30-CA14051 from the National Cancer Institute. This work was supported, in whole or in part, by the Gates Foundation INV-009529. The conclusions and opinions expressed in this work are those of the author(s) alone and shall not be attributed to the Foundation. Under the grant conditions of the Foundation, a Creative Commons Attribution 4.0 License has already been assigned to the Author Accepted Manuscript version that might arise from this submission. Please note works submitted as a preprint have not undergone a peer review process.

All data needed to evaluate the conclusions in the paper are present in the paper and/or the Supplementary Materials.

## Contributions

D.A.S, A.R.K., R.L., and G.T. conceived the concepts of the study. D.A.S., A.R.K., G.N.W., and

D.E.F. performed *in vitro* formulation development and mucin binding studies. D.A.S. and G.N.W. performed *ex vivo* binding studies. D.A.S. performed *in vitro* BLI kinetic measurement. D.A.S., K.I., J.J., and A.P. performed gastrointestinal localization and *in vivo* localized drug delivery studies in rats. J.M. and N.F. performed LCMS chemical analysis for localized drug delivery studies. All authors read and approved the final manuscript.

## Data Availability

The data that support the findings of this study are available from the corresponding author upon reasonable request. Correspondence and material requests should be addressed to Giovanni Traverso.

