## Supporting Information for "Identification and Validation of Small Molecules with Mucin-Selective Regiospecific Binding in the Gastrointestinal Tract"

**Table S1.** Structure and index of 72 selected molecules.

| | | | | | |
|---|---|---|---|---|---|
| **232** | **514** | **517** | **711** | **1001** | **1026** |
| 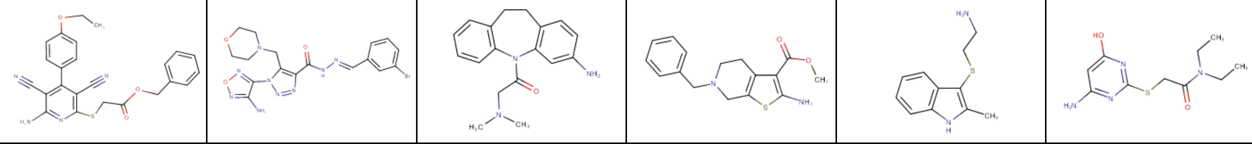 | | | | | |
| **1085** | **1098** | **1206** | **1299** | **1335** | **1336** |
| 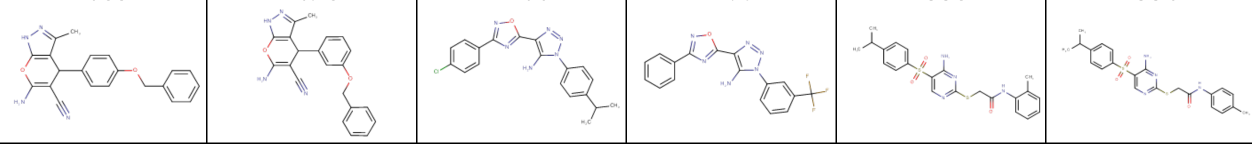 | | | | | |
| **1349** | **1393** | **1394** | **1653** | **1691** | **1855** |
| 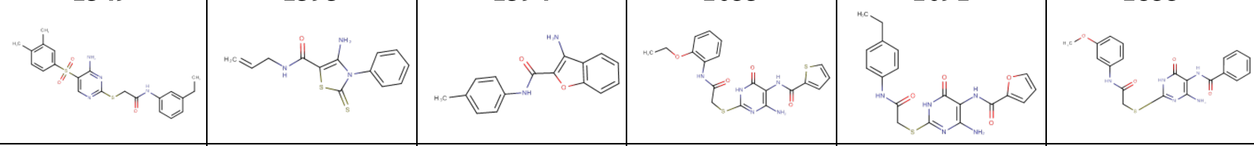 | | | | | |
| **1856** | **1870** | **1978** | **2073** | **2206** | **2235** |
| 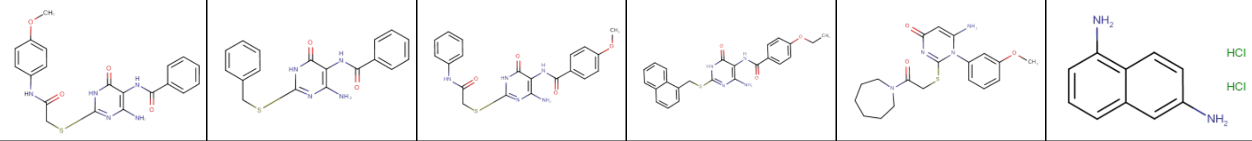 | | | | | |
| **2765** | **2940** | **2951** | **3024** | **3026** | **3031** |
| 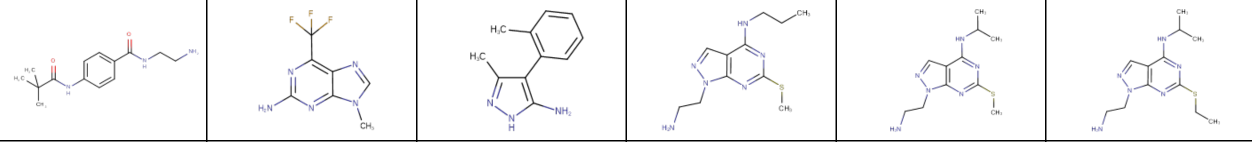 | | | | | |
| **3207** | **3532** | **3576** | **3579** | **3580** | **3609** |
| 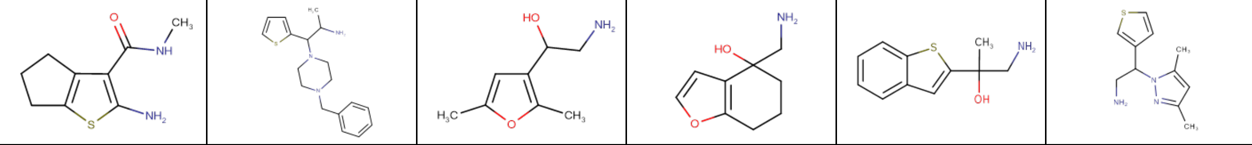 | | | | | |
| **3612** | **3629** | **3640** | **4206** | **4268** | **4273** |
| 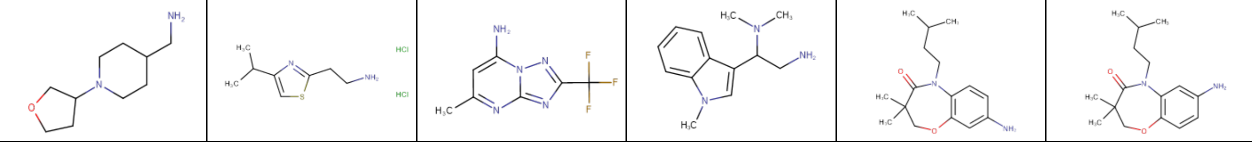 | | | | | |
| **4283** | **4346** | **4353** | **4384** | **4405** | **4426** |
| 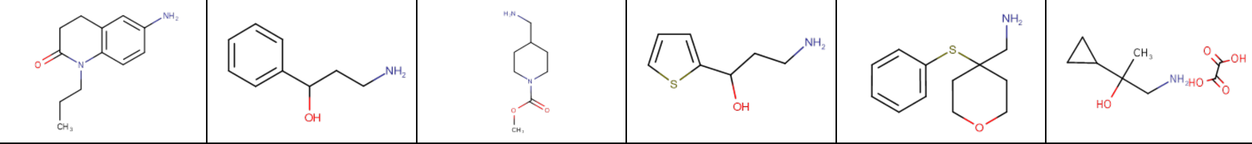 | | | | | |
| **4479** | **4652** | **4664** | **4677** | **4757** | **4961** |
| 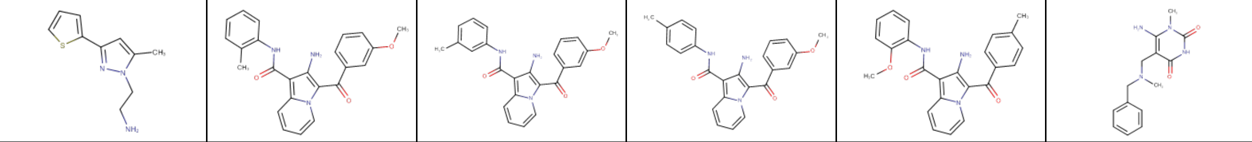 | | | | | |
| **5196** | **5204** | **5398** | **5399** | **6065** | **6070** |

| | | | | | |
|---|---|---|---|---|---|
| 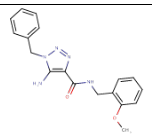 | 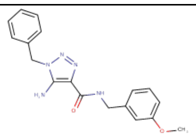 | 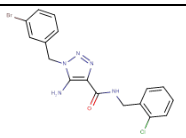 | 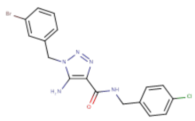 | 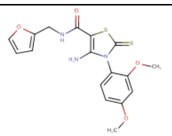 | 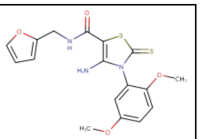 |
| **6074**<br>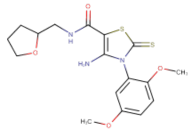 | **6087**<br>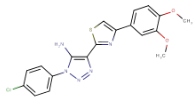 | **6088**<br>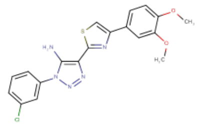 | **6222**<br>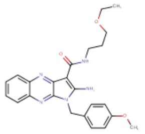 | **6225**<br>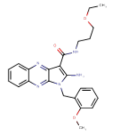 | **6244**<br>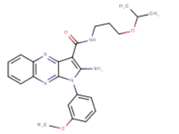 |
| **6626**<br>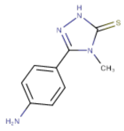 | **6941**<br>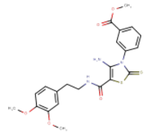 | **7376**<br>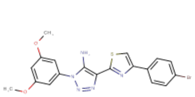 | **7733**<br>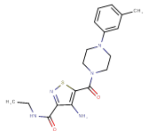 | **8443**<br>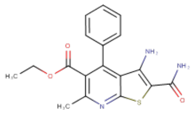 | **8545**<br>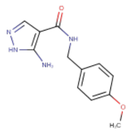 |

**Table S2.** Mucin-binding kinetic constants and dissociation constants for small molecule "hits."

| Mucin | Buffer/pH | Molecule | $k_{on,1}$ $(M^{-1}s^{-1})$ | $k_{off,1}$ $(s^{-1})$ | $k_{on,2}$ $(M^{-1}s^{-1})$ | $k_{off,2}$ $(s^{-1})$ | $k_{d,1}$ (M) | $k_{d,2}$ (M) |
|---|---|---|---|---|---|---|---|---|
| MUC2 | PBS/7.2 | 3207 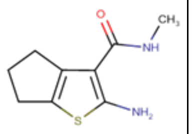 | $1.74 \times 10^{3}$ | $2.75 \times 10^{-}$[illegible] | $3.43 \times 10^{4}$ | $5.65 \times 10^{-}$[illegible] | $1.58 \times 10^{-6}$ | $1.65 \times 10^{-6}$ |
| | | 1026 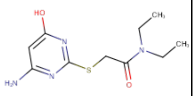 | $2.50 \times 10^{4}$ | $1.53 \times 10^{-}$[illegible] | N/A | N/A | $6.12 \times 10^{-11}$ | N/A |
| | | 1299 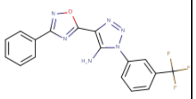 | $1.80 \times 10^{4}$ | $3.95 \times 10^{-}$[illegible] | N/A | N/A | $2.19 \times 10^{-7}$ | N/A |
| | PBS/3.35 | 3207 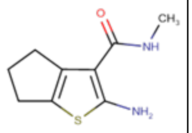 | $7.38 \times 10^{3}$ | $7.46 \times 10^{-}$[illegible] | $6.33 \times 10^{5}$ | $3.06 \times 10^{-}$[illegible] | $1.01 \times 10^{-7}$ | $4.83 \times 10^{-9}$ |
| | | 1026 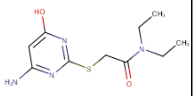 | $3.10 \times 10^{6}$ | $1.45 \times 10^{-}$[illegible] | N/A | N/A | $4.68 \times 10^{-10}$ | N/A |
| | | 1299 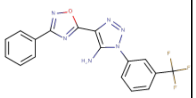 | $2.53 \times 10^{7}$ | $5.51 \times 10^{-}$[illegible] | N/A | N/A | $2.18 \times 10^{-10}$ | N/A |
| MUC5AC | PBS/1.82 | 3207 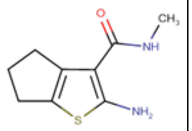 | $3.99 \times 10^{4}$ | $1.49 \times 10^{-}$[illegible] | N/A | N/A | $3.74 \times 10^{-8}$ | N/A |
| | | 1026 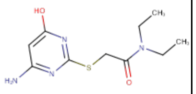 | $6.89 \times 10^{4}$ | $8.62 \times 10^{-}$[illegible] | N/A | N/A | $1.25 \times 10^{-8}$ | N/A |
| | | 1299 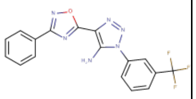 | $1.25 \times 10^{4}$ | $2.47 \times 10^{-}$[illegible] | N/A | N/A | $1.97 \times 10^{-10}$ | N/A |

**Table S3.** Background-subtracted and blank-subtracted fluorescence counts in the esophagus, stomach, and intestines for the top MUC2-binding hits, negative control, and positive control.

| 3207 | | | 1026 | | | N | | | P | | |
|---|---|---|---|---|---|---|---|---|---|---|---|
| Esophagus | Stomach | SI+colon | Esophagus | Stomach | SI+colon | Esophagus | Stomach | SI+colon | Esophagus | Stomach | SI+colon |
| 8.11E+07 | 1.84E+09 | 5.21E+10 | 1.38E+08 | 4.93E+09 | 8.63E+10 | 2.07E+06 | 6.91E+08 | 3.99E+08 | 1.54E+08 | 1.82E+09 | 2.28E+10 |
| 1.02E+08 | 4.04E+09 | 2.09E+10 | 1.76E+08 | 8.98E+09 | 3.49E+10 | 6.94E+06 | 1.03E+09 | 1.34E+10 | 3.57E+07 | 3.88E+09 | 3.76E+10 |
| 1.31E+07 | 8.57E+08 | 1.95E+09 | 3.18E+07 | 2.92E+08 | 7.93E+09 | 1.77E+05 | 2.14E+08 | 1.98E+08 | 1.16E+08 | 6.94E+08 | 3.42E+09 |
| 4.54E+07 | 1.37E+08 | 6.14E+09 | 4.53E+06 | 1.34E+08 | 5.24E+09 | 2.69E+06 | 2.15E+08 | 8.18E+08 | 8.03E+06 | 5.68E+08 | 7.50E+09 |
| 3.40E+08 | 1.51E+09 | 1.37E+10 | 2.99E+08 | 4.25E+09 | 5.06E+10 | 1.10E+07 | 4.06E+08 | 4.84E+09 | 1.01E+08 | 7.06E+08 | 1.08E+10 |
| 7.42E+07 | 4.60E+09 | 2.60E+10 | 1.82E+08 | 3.46E+09 | 3.52E+10 | 2.46E+06 | 6.96E+07 | 5.59E+09 | 1.19E+08 | 1.57E+08 | 8.06E+09 |

**Table S4.** Background-subtracted and blank-subtracted fluorescence counts in the esophagus, stomach, and intestines for the top MUC5AC-binding hit, negative control, and positive control.

| 1299 | | | N | | | P | | |
|---|---|---|---|---|---|---|---|---|
| Esophagus | Stomach | SI+colon | Esophagus | Stomach | SI+colon | Esophagus | Stomach | SI+colon |
| 1.17E+07 | 7.40E+08 | 6.36E+09 | 0.00E+00 | 1.28E+08 | 8.53E+08 | 1.23E+08 | 3.47E+08 | 6.52E+09 |
| 3.44E+07 | 1.05E+09 | 5.67E+09 | 0.00E+00 | 9.40E+07 | 3.95E+08 | 2.08E+07 | 1.20E+08 | 4.38E+09 |
| 1.02E+08 | 4.13E+09 | 1.69E+10 | 0.00E+00 | 1.55E+08 | 2.42E+09 | 6.06E+06 | 5.50E+08 | 6.72E+09 |
| 0.00E+00 | 2.88E+09 | 1.06E+10 | 2.22E+07 | 1.58E+08 | 6.96E+09 | 6.03E+07 | 8.40E+08 | 1.51E+10 |
| 1.95E+06 | 9.91E+08 | 6.62E+09 | 0.00E+00 | 8.93E+07 | 5.36E+08 | 0.00E+00 | 2.46E+08 | 2.68E+09 |
| 0.00E+00 | 7.92E+08 | 2.79E+09 | 0.00E+00 | 1.03E+08 | 3.14E+09 | 8.12E+07 | 4.23E+08 | 6.57E+09 |

**Table S5.** Budesonide fold changes in *in vivo* delivery study versus MUC2-selective hit.

| Stomach | | | SI | | |
|---|---|---|---|---|---|
| 1026 | Unconjugated | Free drug | 1026 | Unconjugated | Free drug |
| 0 | 0 | 0 | 1.94428 | 0 | 0 |
| 0 | 0 | 0 | 1.614649 | 0.480106 | 0 |
| 4 | 0 | 0 | 0.406819 | 0.224377 | 0 |
| 0 | 0 | 0 | 0.034251 | 0.629179 | 0.086658 |

**Table S6.** Mass of tetracycline found in different organs in *in vivo* drug delivery study for MUC5AC-selective hit, negative control, and positive control.

| Stomach | | | SI | | |
|---|---|---|---|---|---|
| **1299** | **Unconjugated** | **Free drug** | **1299** | **Unconjugated** | **Free drug** |
| 103.5381 | 12.94038 | 0 | 25.67962 | 0 | 0 |
| 41.1835 | 0 | 0 | 21.86817 | 0 | 16.16583 |
| 1.344577 | 0 | 0 | 2.108335 | 0 | 0 |
| 58.34617 | 0 | 0 | 35.32762 | 0 | 0 |
| 73.26731 | 0 | 0 | 44.41809 | 0 | 0 |
| 11.5136 | 0 | 0 | 20.81307 | 0 | 0 |
| 4.338248 | 0 | 0 | 9.500837 | 0 | 0 |
| 50.44105 | 0 | 0 | 101.5108 | 0 | 0 |

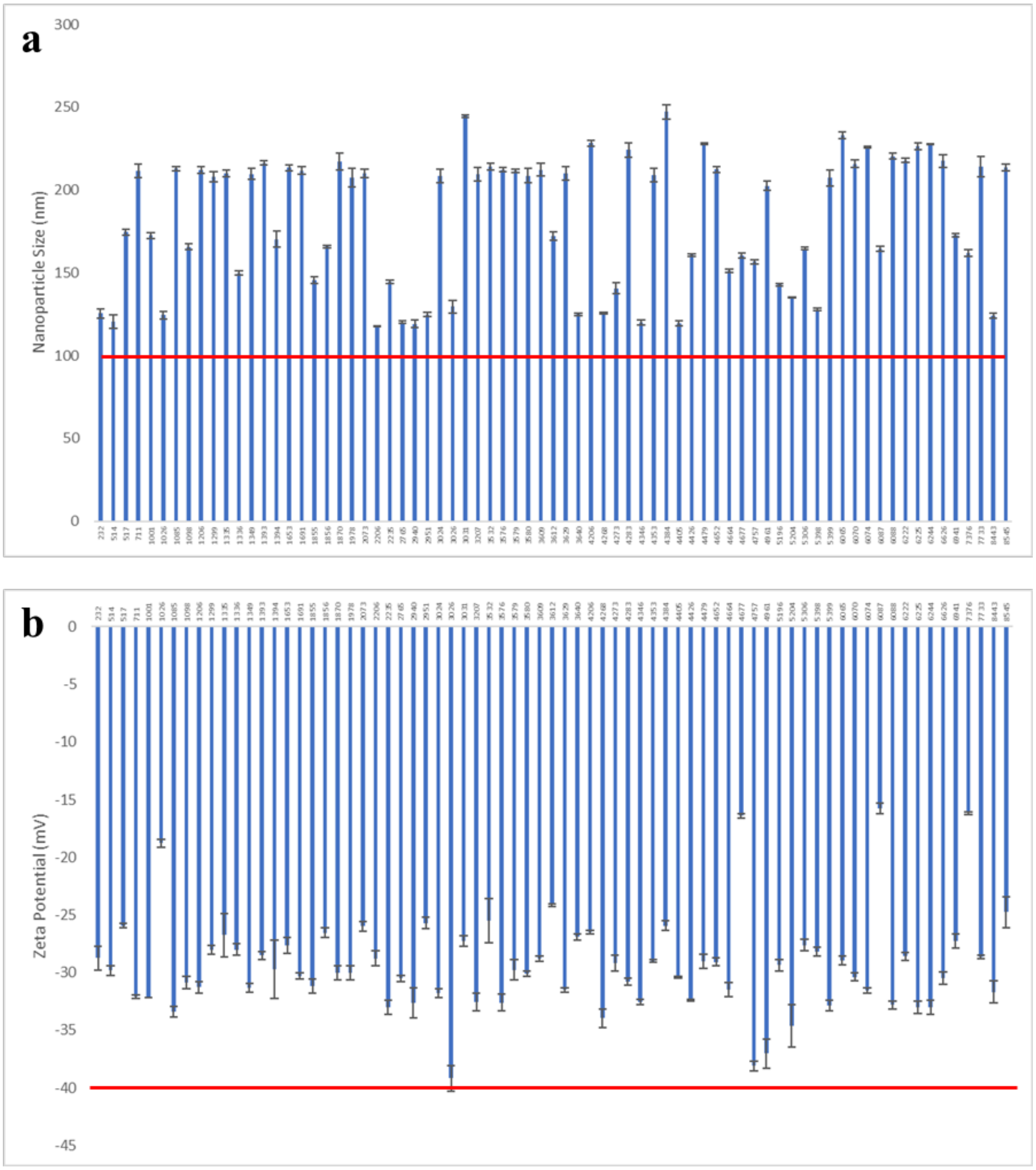


**Figure S1.** Formulation size (a) and zeta potential (b) for the small molecule-conjugated fluorescent nanoparticles. Baseline values for the nanoparticle size and zeta potential for unconjugated nanoparticles are shown with the red lines.

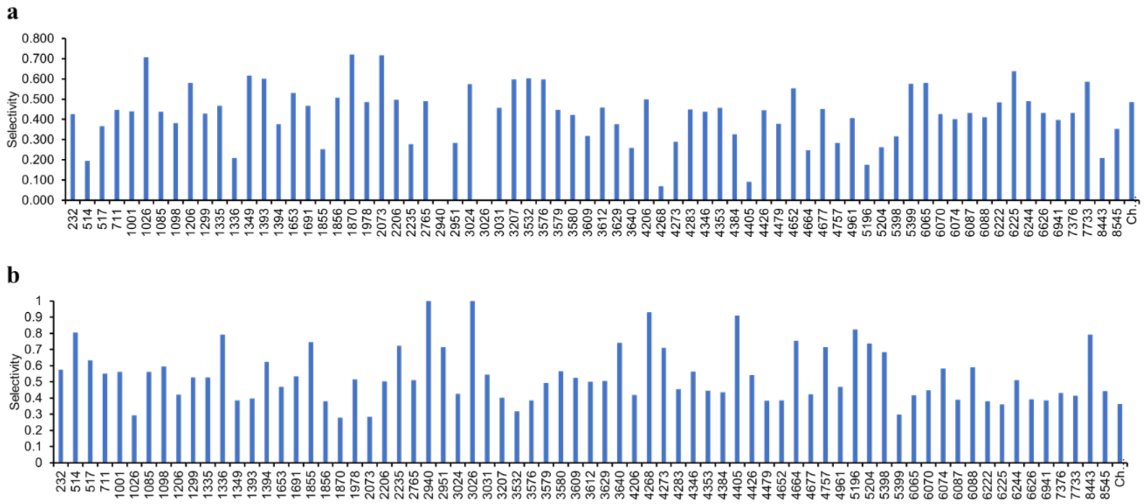


**Figure S2.** Detailed results from the *in vitro* screening of the 72 hits, detailing selectivity to MUC2 (a) and selectivity to MUC5AC (b). Selectivity is calculated by taking the relative contribution of percentage of nanoparticles bound (seen in **Figure 2**) across all mucins. Results represent the median of five replicates.

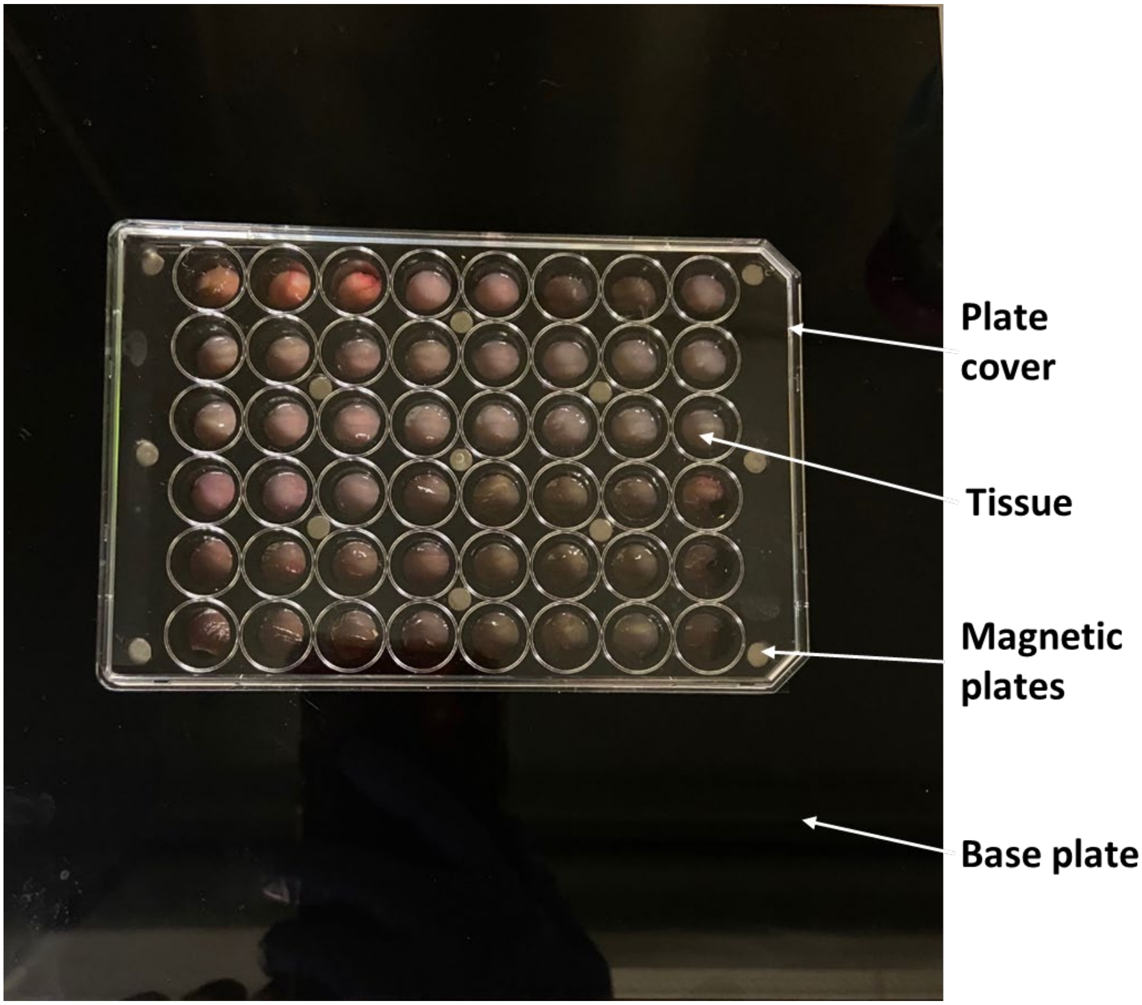


**Figure S3.** Mounted setup for *ex vivo* tissue binding. The setup consists of four parts. The bottom plate is a black polystyrene plate (20 cm x 20 cm), which serves to minimize background fluorescence during imaging studies. The next two plates are magnetic plates with 48 wells cut into the plates. These can suspend a piece of tissue through the magnetic force that holds the plates together, with the luminal side of the tissue facing upwards, allowing for incubation of mucin-binding formulations. The final plate is a standard 48-well plate cover, which is used to enclose the tissue setup.

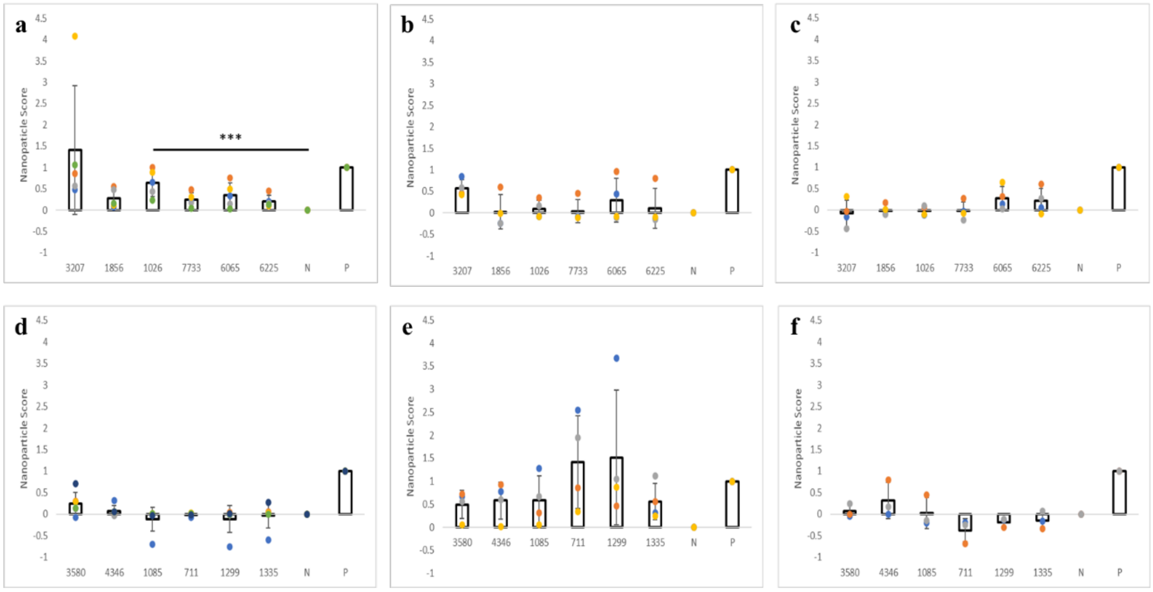


**Figure S4.** Detailed *ex vivo* binding results for MUC2-selective hits to the small intestine (a), stomach (b), and esophagus (c) and MUC5AC-selective hits to the small intestine (d), stomach (e), and esophagus (f). Colored dots represent individual data points. Values on the y-axis correspond to the nanoparticle score, which is defined as $\boldsymbol{Score_i} = \frac{f_i - f_n}{f_p - f_n}$. *** - $p < 0.005$

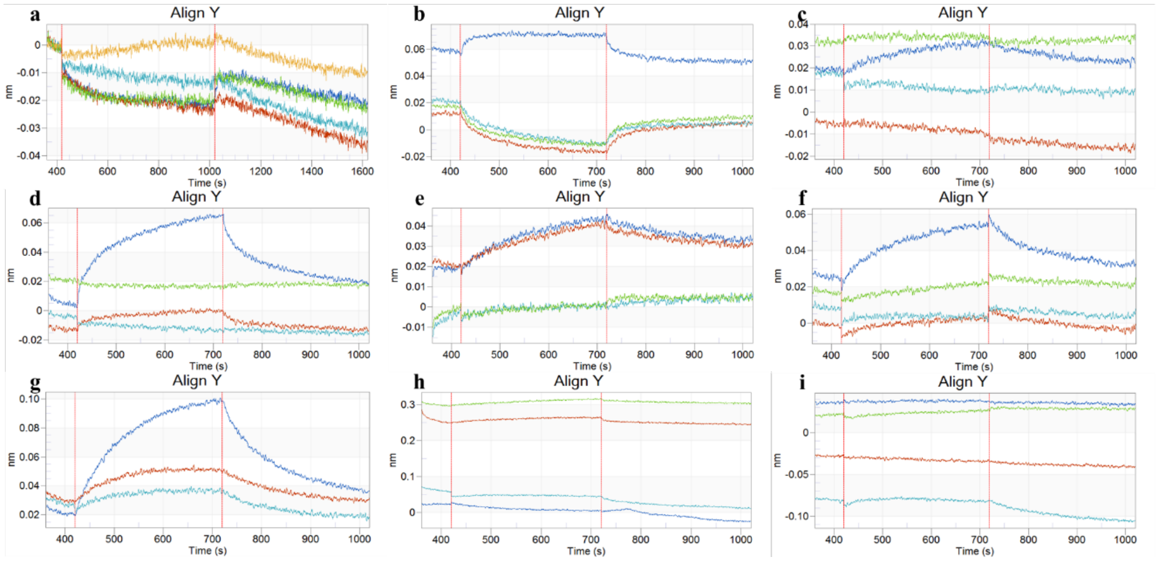


**Figure S5.** Association and dissociation curves measured by bio-layer interferometry. Results shown are for hit 1026 to MUC2 at pH 7.2 (a), MUC2 at pH 3.35 (b), and MUC5AC at pH 1.82 (c), hit 3207 to MUC2 at pH 7.2 (d), MUC2 at pH 3.35 (e), and MUC5AC at pH 1.82 (f), and hit 1299 to MUC2 at pH 7.2 (g), MUC2 at pH 3.35 (h), and MUC5AC at pH 1.82 (i).

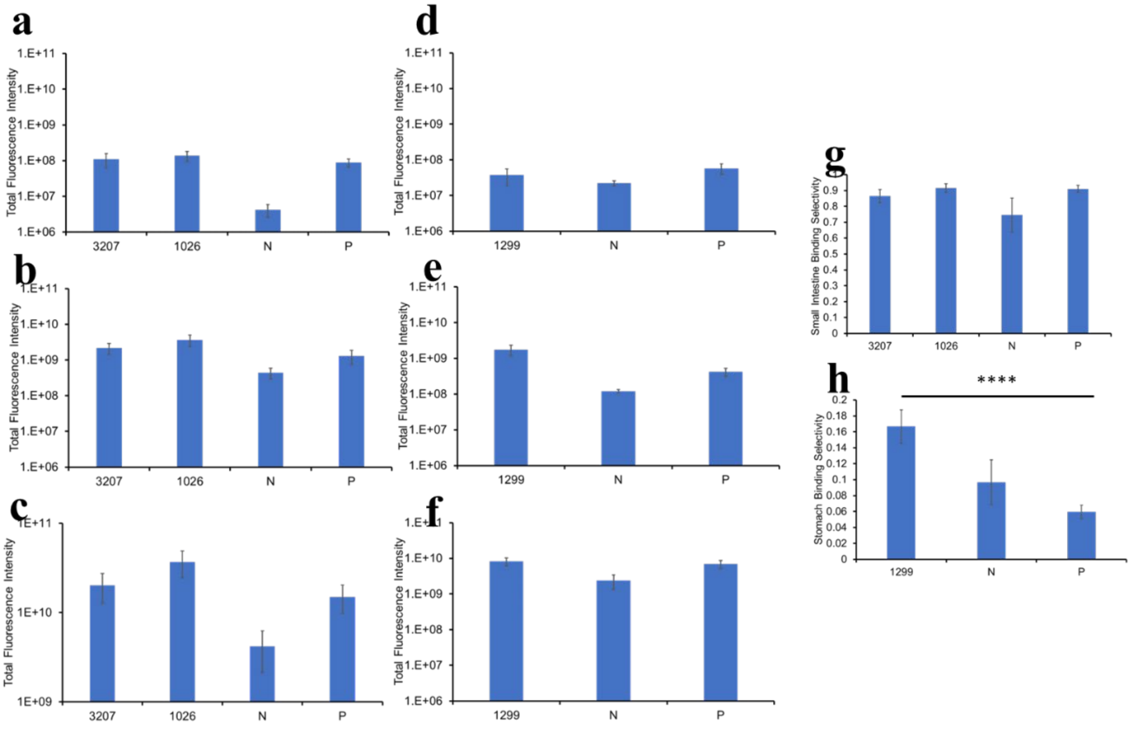


**Figure S6.** Detailed *in vivo* biodistribution results, showing the MUC2-binding hit fluorescence in the esophagus (a), MUC2-binding hit fluorescence in the stomach (b), MUC2-binding hit fluorescence in the small intestine/colon (c), MUC5AC-binding hit fluorescence in the esophagus (d), MUC5AC-binding hit fluorescence in the stomach (e), and MUC5AC-binding hit fluorescence in the small intestine/colon (f). Detailed selectivity results are shown for MUC2-selective (g) and MUC5AC-selective (h) hits. Error bars represent standard error of measurement. **** - $p < 0.0001$.
